# A novel GATA-binding protein 4 gene variation associated with familial atrial septal defect

**DOI:** 10.1101/277780

**Authors:** Yongchao Yang, Yu Xia, Yueheng Wu, Shufang Huang, Yun Teng, Xiaobing Liu, Ping Li, Jimei Chen, Jian Zhuang

## Abstract

Atrial septal defect (ASD) is the most common congenital heart defect. Part of ASD exhibits familial predisposition, but the genetic mechanism remains largely unknown. In the current study, we use multiple methods to identify and confirm the gene associated with a familial ASD. Chromosomal microarray analyses, whole exome sequencing, Sanger sequencing, multiple bioinformatics programs, in silico protein structure modeling and molecular dynamics simulation were performed to predict the pathogenic of the variant gene. Dual-Luciferase reporter gene assay was performed to evaluate the influence of downstream target gene of the target variation. A novel, heterozygous, missense variant GATA-binding protein 4 (*GATA4*):c.958C>T, p.R320W was identified. An autosomal dominant inheritance pattern with incomplete penetrance was observed in the family. Multiple prediction indicate the variant in *GATA4* to be deleterious. Molecular dynamics simulation further revealed that the variation of p.R320W could prevent the zinc finger of *GATA4* from interacting with the DNA. Dual-Luciferase reporter assay demonstrated a significant decrease in transcriptional activity (0.90±0.099 vs 1.50±0.079, *p* = 0.001) of the variant *GATA4* compared with the wild type. We believe the novel variation of *GATA4* (c.958C>T, p.R320W) with a pattern of incomplete inheritance that may be highly associated with this familial ASD. The finding enriched our knowledge of variations that may associated with ASD.

## Introduction

Congenital heart defect (CHD) is one of the most common life-threatening birth defects, with an estimated incidence of 7 to 9 per one thousand and affecting 100,000 to 150,000 newborns in China each year.^1-3^ There are more than 400 identified variant genes, including *NKX2-5, MYH6, GATA4, ZIC3*, and *ELN*, associated with CHD and accounting for about 10% of CHD.^4-7^ The ostium secundum atrial septal defect (ASD), comprising of approximately 30 to 40% of CHD, is one of the most common subtypes of interatrial communication,^8^ and often associated with other cardiac and/or extra-cardiac anomalies and genetic syndromes.^9^

The understanding of the genetic pathogenesis mechanisms of ASD has significantly improved in recent decades. Pathogenic genomic copy number variants (CNVs) and gene variants have been identified in ASD. Rare CNVs at recurrent loci, such as 22q11.2, 5q35.1, 8p23.1, and 18q11.2, and more than ten genes, including as *NKX2-5, GATA4, GATA6, TBX5, TBX20*, and *CITED2*, have been associated with sporadic or inherited ASD.^10-12^

*GATA4*, a zinc finger transcription factor that contains seven exons located on chromosome 8p23.1-p22, plays an important role in early stage of embryonic heart development.^13,14^ It is comprising of 442 amino acids with four conserved domains – transcription activation domain 1(TAD1, amino acids (aa) 1 to 74), transcription activation domain 2 (TAD2, aa 130 to 177), N-terminal zinc finger (ZF1, aa 217 to 241), and C-terminal zinc finger (ZF2, aa 271 to 295).^14,15^ The nuclear localization signal (NLS, aa 271 to 325) allows GATA4 to be imported to the nucleus via the nuclear pore complex.^14^ Small changes in the level of GATA4 protein expression can dramatically influence cardiac development and embryonic survival. Presently, there are 150 variant sites of *GATA4* that have been identified in association with different subtypes of CHD, including atrial septal defects, ventricular septal defects, tetralogy of Fallot, and atrial fibrillation.^16-19^ There are 104 missense or nonsense variants of *GATA4* reported to be associated with these CHD in Human Gene Mutation Database (HGMD) (http://www.hgmd.cf.ac.uk/ac/all.php).

However, as a genetically heterogeneous disease, such a large number of variations still cannot fully cover the variations of ASD. Here, we use multiple methods to identify and confirm the variant gene associated with a familial ASD.

## Materials and methods

### Study subjects

A family with 8 members were enrolled in this study (II-2, III-1, III-2, III-3, IV-3, IV-4, IV-5 and IV-6, Figure 1). All enrolled members underwent a complete physical examination. Clinical data, including medical records, electrocardiograms, and echocardiography were systematically reviewed. The study protocol (protocol number 20140829) was approved by the Research Ethics Committee of Guangdong General Hospital, Guangdong Academy of Medical Sciences, Guangdong, China. Informed written consent (informed consent form number 20150424) was obtained from all family members. Approximately 6.0 mL of peripheral blood was collected from each of the study participants. DNA was extracted from the peripheral blood lymphocytes using modified salting-out precipitation method by Gentra Puregene blood kit (QIAGEN, Santa Clara, CA, USA).

**Figure 1.**
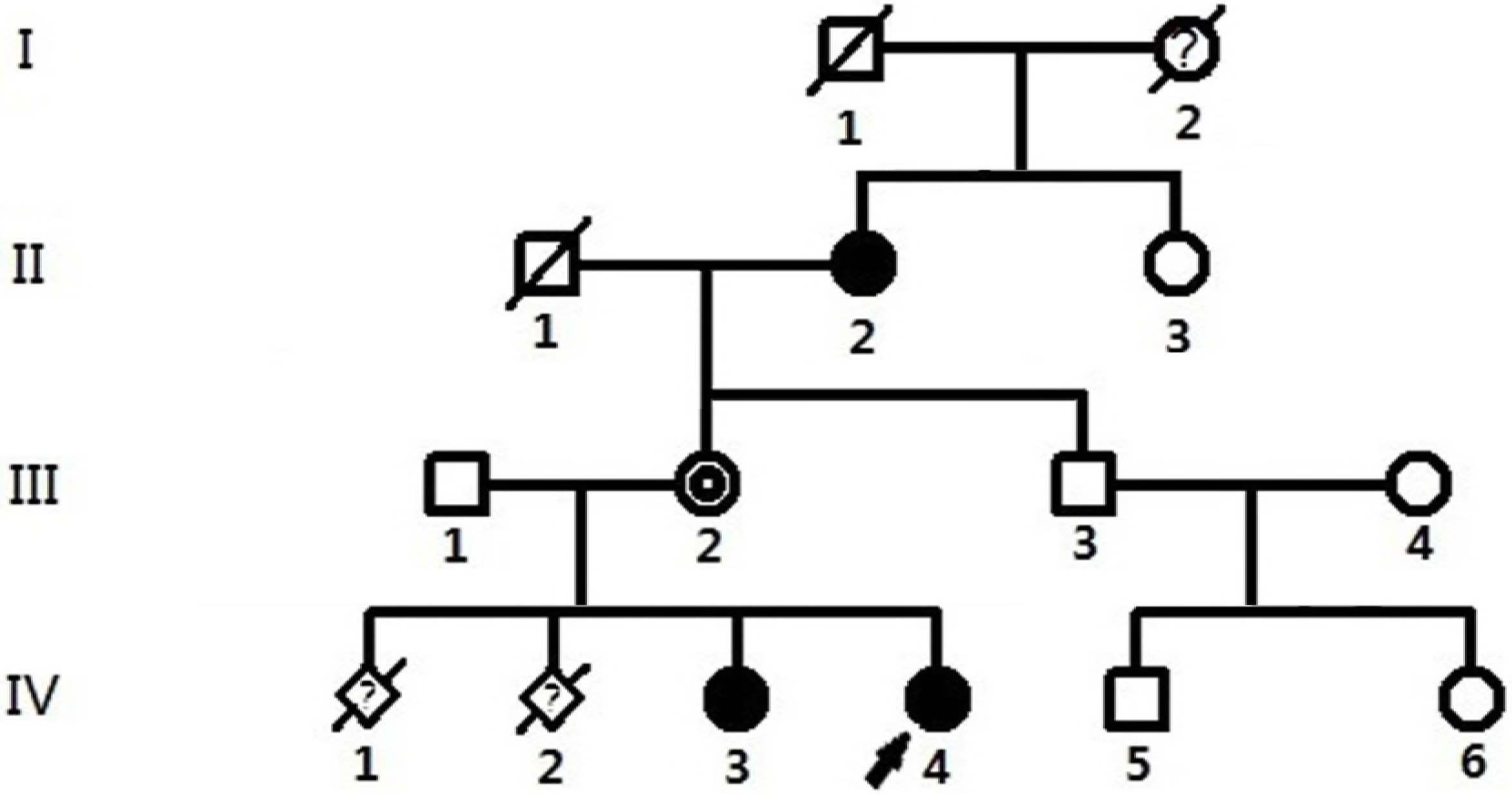
Pedigree of the nuclear and extended family showing affected and unaffected members with ASD. Subjects I1 and II1 died naturally. Subject I2 was a suspected positive. Subjects II2, IV3, and IV4 were affected patients. Subject III2 was a carrier. Subject IV1, miscarried at the 8th week of pregnancy. Subject IV2, mid-trimester aborted for fetal bradycardia. The arrow indicates the proband patient. ‘?’ indicates the suspected patient. The oblique line represents a deceased subject.

### Chromosomal microarray analysis and CNV evaluation and validation

Two hundred fifty nanograms (ng) of DNA was amplified from III-1, III-2, IV-3, and IV-4 (Fig. 1), then labeled and hybridized to the CytoScan HD array platform (Affymetrix, USA) according to the manufacturer’s protocol. The array was designed specifically for cytogenetic research, offering more than 2,700,000 markers across the whole genome, including 750,000 SNP probes and 1,950,000 probes to detect CNVs (Cyto-arrays). Data were visualized and analyzed with the Chromosome Analysis Suite (ChAS) software package (Affymetrix, USA) with a minimal cutoff of 20 consecutive markers in a length of 25-kb for CNVs calling. All of the segments were monitored for the degree of overlap with previously identified common CNVs, annotated by Database of Genomic Variants (DGV). All of the reported CNVs are based on NCBI human genome build 37 (hg 19).

Detected CNVs meeting the following criteria were selected for further analysis: (1) deletions greater than or equal to 50 kb and duplications greater than or equal to 50 kb; (2) without recurrence in the normal populations that have been cataloged in DGV; and (3) possessing less than 50% overlap with known segmental duplications.

Following the American College of Medical Genetics and Genomics’ (ACMG) standards and guidelines for the interpretation of CNVs, the remaining CNVs were classified into three categories—pathogenic (P), variants of uncertain significance (VOUS), and benign (B). VOUS was further divided into three parts—likely pathogenic (LP), likely benign (LB), and no sub-classification (NS). For this study, only genes that function in a dominant manner that are within P and LP CNVs were investigated. All of the annotated CNVs were experimentally validated by real-time quantitative PCR (qPCR).

### Whole exome sequencing and variant analysis

To systematically search for disease-causing genes, exome sequencing in three affected individuals and two unaffected individuals (parents of proband) from the family with a history of ASD was performed using the Agilent Sure Select Human All Exon V5 Kit on the Illumina HiSeq 2000 platform by Novogene Bioinformatics Technology Co., Ltd. One and 0.5 µg of genomic DNA from the proband was used to construct the exome library. The genomic DNA was sheared into fragments with a length of 180 to 280 bp by sonication and hybridized for enrichment according to the manufacturer’s protocol. The library enriched for target regions was sequenced on the Illumina HiSeq 2000 platform to get paired-end reads with a read length of 100 bp. The average sequencing depth of 57.36X provided enough depth to exactly call variants at 97.4% of the targeted exome.

The human reference genome was obtained from the University of California Santa Cruz (UCSC) database (build 37.1, version hg19, http://genome.ucsc.edu/), and sequence alignment was performed using the Burrows-Wheeler Alignment tool. High-quality alignment was required to guarantee variant calling accuracy (greater than 0). Picard (http://sourceforge.net/projects/picard/) was employed to mark duplicates resulting from PCR amplification. Genome Analysis Toolkit (GATK) Indel Realigner and GATK Realigner Target Creator were performed to do realignment around the indels. GATK Base Recalibrator was performed to do base quality score recalibration. GATK Variant Filtration was performed to make the raw callsets suitable for meaningful analysis. Exome CNV was performed for CNV detection.

Sequence Alignment/Map (SAM) tools were used to perform variant calling and identify single nucleotide polymorphisms (SNPs) or indels. After the analysis-ready Binary Alignment/Map (BAM) alignment result was obtained, Annotate Variation (ANNOVAR) was performed to annotate SNPs and indels. All candidate variants were filtered against the Single Nucleotide Polymorphism Database (dbSNP142, http://www.ncbi.nlm.nih.gov/projects/SNP/snp_summary.cgi), 1000 Genomes Project (2016 April release, http://www.1000genomes.org/), Exome Aggregation Consortium (ExAC, http://exac.broadinstitute.org/) and NHLBI Exome Sequencing Project (ESP) 6500 to remove the polymorphism loci. Sorting Intolerant from Tolerant (SIFT) and Polymorphism Phenotyping version 2 (PolyPhen-2), Mutation Taster, and Combined Annotation Dependent Depletion (CADD) were performed to predict whether an amino acid substitution affects the function of the protein.

In addition to the standard variant quality controls, six independent filters were applied to facilitate detection of possible causal variants among the enrolled ASD patients. Variants were filtered by: (1) a minor allele frequency (MAF) of less than 1% in east Asian population; (2) CADD score of greater than 20; (3) the variants must be pathogenic; (4) highly expressed in the heart or associated with cardiac development; (5) genotype–phenotype matched under the assumption of complete penetrance.

### Sanger sequencing confirmation

Direct Sanger sequencing was performed with ABI 3500 sequencer (Applied Biosystems, Foster City, CA, USA) to confirm potential causative variants in the family. Primer sequences for pathogenic variant in the *GATA4* gene (NM_002052.4) were designed as follows: 5’-CAATGCCTGCGGCCTCTAC-3’ and 5’-AGGAAGAAGACAAGGGAGGACTG-3’. Once a variant was confirmed, all family members were screened to analyze variant segregation within the family.

### Conservation analysis across species, in silico protein structure modeling and molecular dynamics simulation

The protein sequences of GATA4 in 11 species from *Drosophila melanogaster* to *Homo sapiens* were aligned using Clustal X (version 1.81) software.

The wild type GATA4 protein from the previous homology modeling study was used as the initial structure in molecular dynamics (MD) simulations. Then, the energy minimized and equilibrated structure was used to acquire the R320W mutant protein by using UCSF Chimera.^20^ Finally, 10 ns MD simulations were performed for both wild type and mutant proteins of GATA4. All preparation and simulations were performed in AmberTools15 (Amber 2015, University of California, San Francisco). The Leap module was employed to assign AMBER ff14SB force field for protein and zinc (Zn^2+^) ions. Zn^2+^ ions with coordinate atoms were constrained to be tetrahedron using the cationic dummy atom (CaDA) approach of Pang et al.^21^ TIP3P water model was used and the box was set to 10 Å. Counter ions Cl^-^ were added in order to neutralize charges of the system. A 30 ns of NPT ensemble MD simulation was performed using Lengevin dynamics method^22^ to control temperature with collision frequency of 1.0 ps^-1^. A SHAKE algorithm was applied to constrain bonds involving hydrogen atoms. The van der Waals cutoff was kept 10 Å and long range electrostatic interactions were treated using the Particle Mesh Ewald (PME)^23^ method. Atomic coordinates were saved after every 500 steps. RMSD, RMSF, radial gyration (Rg), solvent accessible surface area (SASA) and secondary structure analyses were carried out to study the effect that mutagenesis would have on the structure and functions of GATA4 protein.

### Plasmids and site-directed mutagenesis

The full-length wild type cDNA of the human *GATA4* gene was amplified by PCR using PrimeSTAR^®^ HS DNA Polymerase (Takara, Liaoning, CHN) and primers (5’-GGGGTACCATGTATCAGAGCTTGGCCATGGCC-3’ and 5’-CCGCTCGAGTTACGCAGTGATTATGTCCCC-GTGA-3’). PCR fragments were double digested by endonucleases Kpnl and Xhol (Thermo Fisher, Shanghai, CHN). The digested product was fractionated by using 1.5% agarose gel electrophoresis, purified by using the E.Z.N.A^®^ Gel Extraction Kit (OMEGA, Norcross, GA, USA), and then subcloned into pcDNA3.1 (Promega, Beijing, CHN) to construct the recombinant eukaryotic expression vector WT-pcDNA3.1-hGATA4.

The *GATA4* variant c.958C>T (p.R320W) was introduced into a wildtype *GATA4* clone using a QuikChange Site-Directed Mutagenesis Kit (Stratagene, Agilent Technologies, CA, USA) and primers (5’-ATCCAAACCAGAAAATGGAAGCCCAAGAACC-3’ and 5’-GGTTCTTGGGCTTCCATTTTCTGGTTTGGAT-3’). The clones were sequenced to confirm the expected variant and exclude other variants.

### Dual-Luciferase assays

293T cells were transiently transfected with 400 ng brain natriuretic peptide (*BNP*)-luciferase reporter plasmid and internal control reporter plasmid pGL4.75 [hRluc/CMV] (Promega, Southampton, UK) in combination with 100 ng of wild type *GATA4*, mutant type *GATA4* p.R320W, or pcDNA3 plasmids using Lipofectamine™ 2000 (Invitrogen, Cat.) according to the manufacturer’s protocol. Luciferase activity was measured 48 hours after transfection using the Dual-Glo Luciferase Assay (Promega, Southampton, UK) according to the manufacturer’s protocol. Mean luciferase activity was calculated after normalization to Renilla luciferase activity. The experiments were performed and repeated at least three times respectively. Finally, the expression level of protein GATA4 were evaluated by western blot analysis.

### Statistical Analysis

Statistical analysis was performed with SAS-PC (9.3v, SAS Institute, Inc, Cary, NC). Results are expressed as mean ± Standard Deviation (SD). Logarithmic transformation was applied to the data for the luciferase assay to achieve approximate normality. Comparisons between two groups were performed with the chi-square test. Statistical significance was determined by two-way analysis of variance and subsequent pairwise comparisons were performed. A *p* value of less than 0.05 was considered to be statistically significant.

### Data availability

The authors state that all data necessary for confirming the conclusions presented in the article are represented fully within the article.

## Results

### Clinical Characteristics

The proband of this family is the second child, who was diagnosed with ostium secundum atrial septal defect by ultrasound (IV-4) (Fig. 1). After reviewing the family history, we identified the first child (IV-3) (Fig. 1) was also affected by the same defect and had been cured by transcatheter closure four years prior to our interview. The mother of proband was not detected with any cardiac disorders by echocardiogram (III-2) (Figure 1). The maternal grandmother (II-2) (Fig. 1) indicated upon interview that she had undergone cardiac surgery due to ASD when she was approximately 20 years of age (the detail medical records were missing). She also informed us that her mother (I-2) (Fig. 1) may have had a cardiac disorder as she was unable to perform manual labor and died due to an unidentified cause at approximately 40 years of age. The mother of the proband had two abnormal pregnancies with a spontaneous miscarriage at approximately 8 weeks during the first pregnancy (IV-1) (Fig. 1) and a termination due to the detection of fetal bradycardia at 18^th^ week during the second pregnancy (IV-2) (Fig. 1). Conventional G-banded cytogenetic analysis was performed for patients II-2, III-1, III.2, IV-3 and IV-4, but no clinically significant result was found.

### No significant CNV was found associated with ASD

CMA was performed on DNA samples from members III-1, III-2, IV-3, and IV-4. There was only one subject (III-1) with detection of a 1.641 Mb duplication at Yq11.223 (chrY: 24,148,853-25,790,030). This duplication only contents one disease causing gene of *DAZ1* according to OMIM. The deletion of this gene may be related to spermatogenesis, but it is unknown if its duplication has any significant clinical annotation.

### *GATA4* was located as target variant gene for this familial ASD

Exome analysis was performed on five DNA samples from members II-2, III-1, III-2, IV-3 and IV-4. A total of 19,816,364 pairs of sequenced reads with the average read length of 125 bp were generated by exome sequencing. Approximately 98.76% (19,569,878) of sequenced reads passed the quality assessment and were mapped to 99.81% of the human reference genome. There were more than 20,000 indels, 167,000 SNPs and 10 CNVs found in each subject. After filtering, more than 1500 variants including SNP and indels were shared by II-2, IV-3 and IV-4, and finally, a heterozygous missense variant *GATA4*: NM_002052: exon4: c.C958T: p.R320W (CADD_Phred score: 35, SIFT score: 0.0, Polyphen2_HVAR score: 0.999, Polyphen2_HDIV score: 1.0, Mutation Taster predict: Disease causing), was selected as our target pathogenic variant detected in IV-3, IV-4, III-2 and II-2.

In order to validate the target variant, we performed the Sanger sequencing in all relevant subjects (II-2, III-1, III-2, III-3, IV-3, IV-4, IV-5 and IV-6) (Fig. 2A). The heterozygous *GATA4* p.R320W variant was found in subjects II-2, III.2, IV-3 and IV-4 but not in subjects III-1, III-3, IV-5 and IV-6 (Table 1).

**Figure 2.**
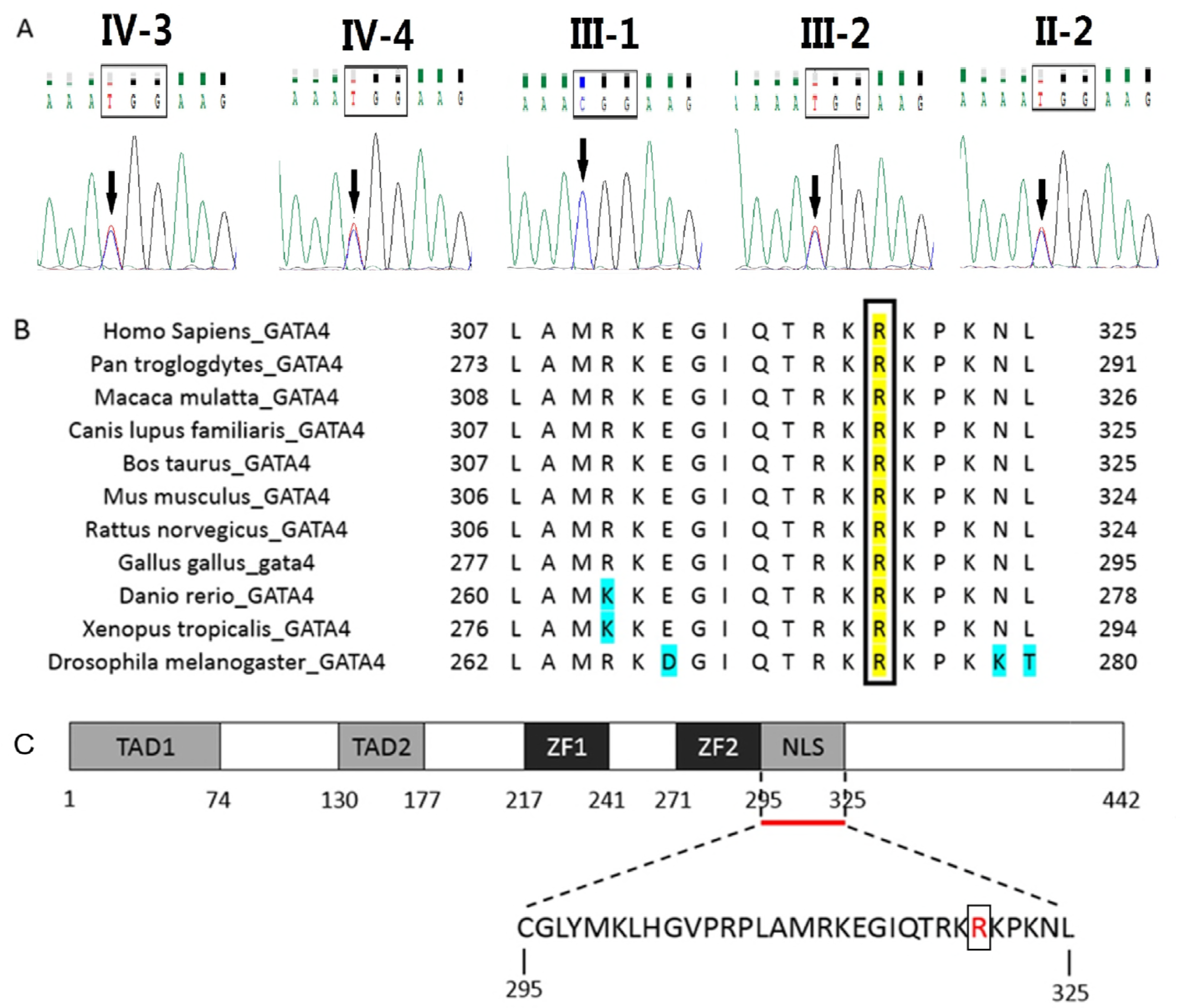
Heterozygous missense variation of *GATA4* in the family and the conservation analysis of the variant site. (A): The arrow indicates the heterozygous nucleotides of C/T. Subject III-1 shows the normal individual. Subjects IV-3, IV-4, III-2 and II-2 show the missense variant. The rectangle denotes the nucleotides comprising a codon of *GATA4*. (B) Alignment of multiple GATA4 amino acid sequences across species. The altered arginine at amino acid position 320 (p.R320) of GATA4 is completely conserved evolutionarily among various species. The yellow column shows the R320 site. The fluorescent blue shows unconserved sites in different species around the R320 site. (C) Schematic diagram of GATA4 protein. TAD1: transcription activation domain 1 (amino acid 1-74); TAD2: transcription activation domain 2 (aa 130–177); ZF1: N-terminal zinc finger (aa 217– 241); ZF2: C-terminal zinc finger (aa 271–295); NLS: nuclear localization signal region (aa 307–325). Red bar shows NLS and the red letters are mutant amino acids (R320>W).

### The variant site of *GATA4* is evolutionarily conserved and plays an important role in the region of nuclear localization signal

A cross-species alignment of the *GATA4* amino acid sequences revealed that the altered amino acid arginine (CGG) at position 320 was completely conserved evolutionarily (Fig. 2B). The identified variant is located in the region of the nuclear localization signal (Fig. 2C), which plays an important role in the nuclear translocation of *GATA4*.^14^

After structure preparation and MD simulation, we found the amino acid structure of *GATA4* in position of 214-322 mainly contains two zinc binding regions linked by a random coil, with the variant site located at one end of coil. In wild type *GATA4*, the random coil is far from the other zinc binding region without any connection. The structure has an “open loop”, with an angle of 167° between the two-helix structures of zinc binding regions (Fig. 3A). In the mutant *GATA4*, connections via hydrogen bonds and hydrophobic interactions between the random coil and the opposite side of zinc binding region were formed. The structure has a “closed loop”, with an angle of about 73° (Fig. 3C). This was likely due to the change of electric charge from the variation of Arg320 to Trp320 where Trp320 formed the hydrophobic interaction with Leu227 and hydrogen bonds with Arg230. The variant also forced a change of the random coil in segments of 256 to 259 into a helical structure (Fig. 3B, 3D).

**Figure 3.**
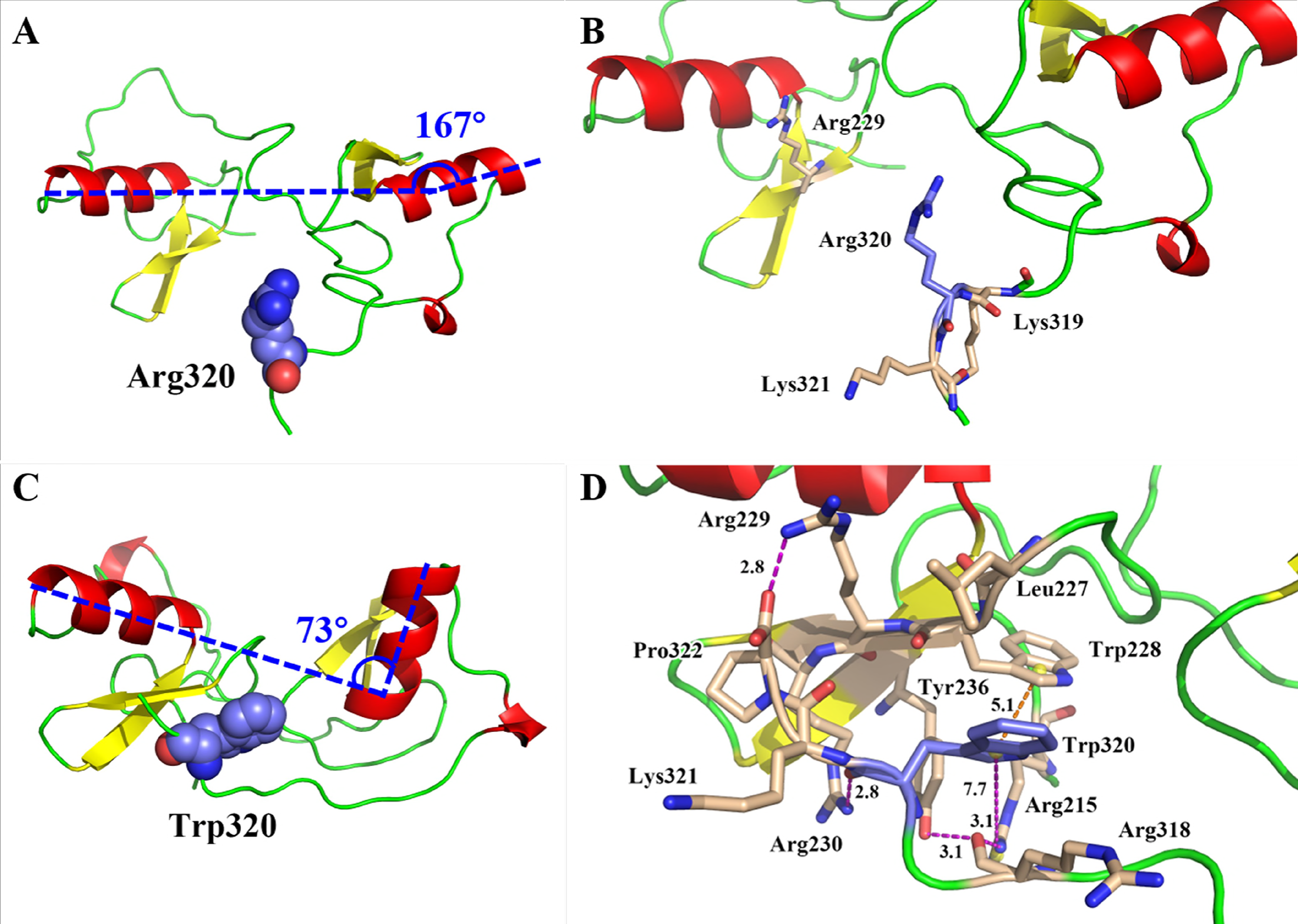
The structural difference between wild type and mutant *GATA4*. (A and B): wild type; (C and D): mutant GATA4. Colorful cartoon depicting the protein of GATA4. Red indicates the α-Helix while yellow indicates the β-Sheet. Green shows a random coil. Sphere and stick of blue shows the variant site. Wheat shows the related residues. Dotted line in (D) shows the new hydrogen bonds. Blue dotted line in (A) and (C) shows the angle of two α-helices.

### Variation of *GATA4* significantly decrease the expression of *BNP*

Previous studies have revealed that *GATA4* is an upstream transcriptional regulator of several genes expressed in different signaling pathway during cardiac development, including genes that encode atrial natriuretic peptide (*ANP*), brain natriuretic peptide (*BNP*), and β-myosin heavy chain (*MHC*).^13^ Therefore, the functional effect of the *GATA4* variant may be reflected by biochemical analysis of the transcriptional activity of the *BNP* promoter in cells transfected with mutant *GATA4* in contrast to its wild type counterpart.

In the dual-luciferase reporter assay (Fig. 4A), we found wild type *GATA4* can significantly increase the transcription of *BNP* in comparison with nature control type (1.50 ± 0.079 vs 1.0 ± 0.064, *p* = 0.001), which in accordance to previous knowledge that *GATA4* is activating transcription factor for *BNP*. The mutant *GATA4* displayed a significant decrease in transcriptional activity in comparison to wild type (0.90 ±0.099 vs 1.50 ± 0.079, *p* = 0.001). There is no significantly difference in protein levels between wild type and mutant GATA4 by analyzing of Western blot (Fig. 4B).

**Figure 4.**
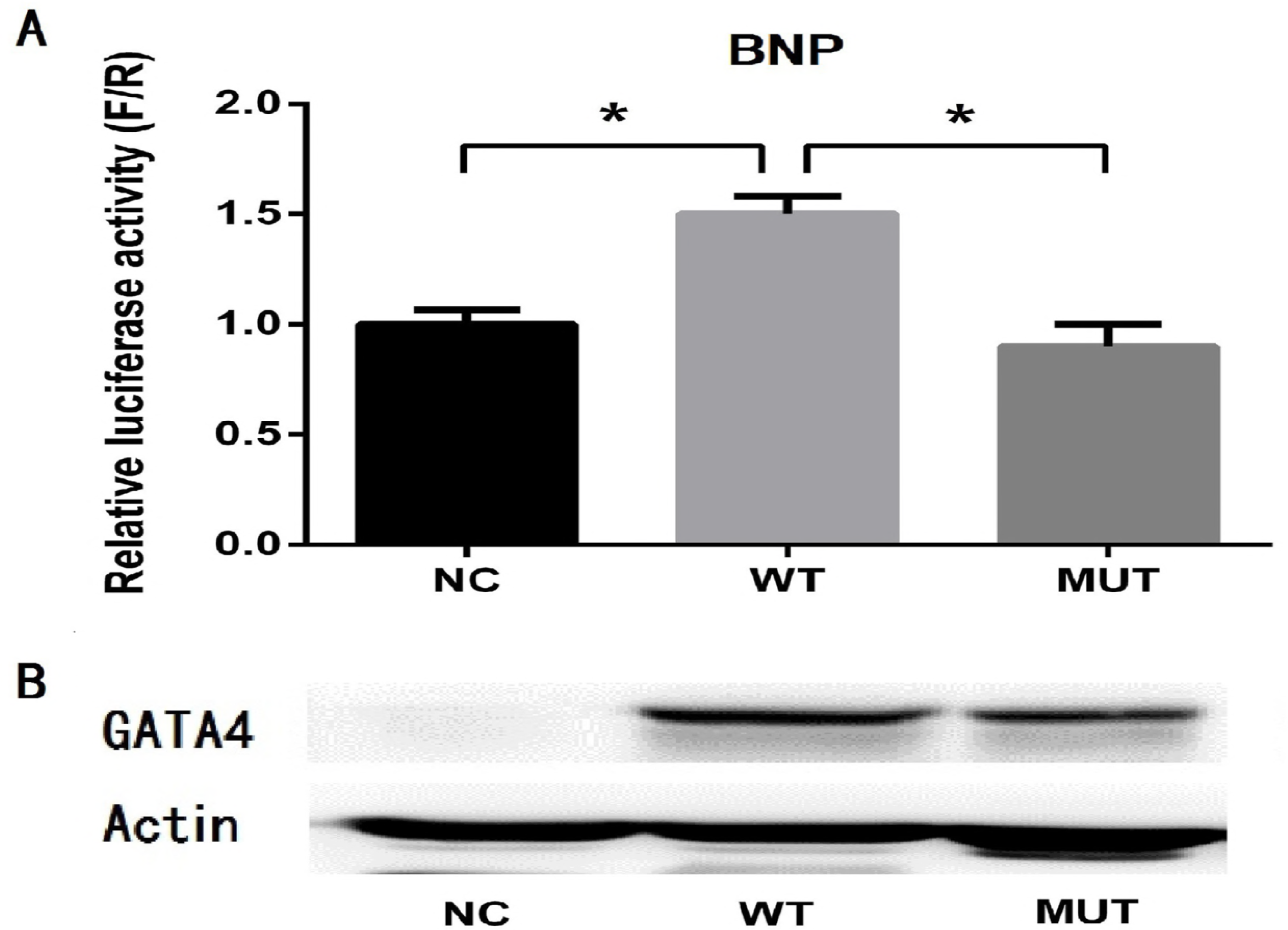
Diminished transcriptional activity of *GATA4* caused by the variation. (A) 293T cells were transfected with 100 ng of wild type, mutant GATA4 or pcDNA3 plasmids and 400 ng of BNP luciferase reporter. The result showed the transcriptional activity of mutant *GATA4* significantly decreased in comparison to wild type. NC: negative control; WT: wild type; MUT: mutant. Data is displayed as mean ± SD; NC was set as 1.0; * represent a statistically significant difference with *p* < 0.01. (B) The protein levels of wild type and mutant GATA4 were evaluated by western blot analysis after 293T cells were transfected with 100 ng of either type of plasmid. No significantly differences were observed between the two groups.

## Discussion

We interviewed and evaluated members of a family with at least 3 patients diagnosed with ASD in the clinic. After a complete examination including karyotyping, CMA, WES and Sanger sequencing for the whole family, we found a novel, heterozygous, missense variation of *GATA4*, c.C958T:p.R320W, in 3 patients with ASD and one unaffected carrier (the mother). This finding was not consistent with traditional autosomal dominant Mendelian inheritance. Interestingly, E. D’Amato et al^24^ reported a heterozygous missense variant *GATA4* c.1512C>T, p.Arg319Trp by HGMD in two children from an Italian family with pancreatic agenesis and ASD, also with the inheritance of incomplete penetrance. There is no completely reasonable explanation for this type of inheritance, it may also be related to complicated gene–gene or gene– environment interactions. With the childbearing history of two terminal pregnancies, the pleiotropism of the gene may explain why the proband’s mother carried the same *GATA4* variant but had distinct clinical phenotypes. This phenomenon may highlight the fact that even in a familial CHD, the underlying genetic etiology can be complex.

There is no record of this variant in the databases of 1000G, ESP6500, ExAC and NCBI. However, the bioinformatics programs across all prediction algorithms, including PolyPhen-2, SIFT, Mutation Taster, and CADD, etc. According to the conclusions of Philips AS, our finding of the *GATA4* variant p.R320W may be pathogenic.^14^ The algorithms demonstrated that four amino acids, Arg282, Arg283, Arg317, Arg319, played crucial roles in nuclear localization of *Gata4* in murine cell line models. Coincidentally, we identified the same variant site of Arg320 (the same site with murine Arg319) in this family. Given that this variant site is highly evolutionarily conserved across 11 species, the variant may lead to a pathogenic change of function after translated into protein. Our prediction of molecular architecture of *GATA4* protein shows the influence of the variant on changing the stability of conformation and structure, which may decrease or inhibit its function in transcription. Therefore, it is very likely that dysfunctional *GATA4* contributes to ASD in this family.

As a key transcription factor, *GATA4* regulates transcription of many genes involved heart development, including *MHC, ANP, BNP* and endothelial nitric oxide synthesis (*eNOS*),^13.25^ and *GATA4* also plays an essential role in cardiac adaptive responses, including myocyte survival, angiogenesis, and hypertrophy in response to exercise.^13,25-27^ In our dual-luciferase reporter assay, we also confirmed that *GATA4* is a transcriptional factor of BNP. The variation of *GATA4* p.R320W significantly decreased its transcriptional activity on downstream cardiac genes of *BNP* in 293T cells, indicating that this variant could contribute to the pathogenesis of ASD.

Our study demonstrates the fact that CMA and WES are reliable techniques in detection of pathogenic variants in familial ASD. Detailed clinical genetic data will facilitate genetic diagnosis and counseling when evaluating the prognosis of newborns and the future risks for other family members with new pregnancies. Our future work will focus on exploring the change of relevant signaling pathway caused by the variation of *GATA4* and the prevalence of this variant site in population of ASD.

## Conclusion

In conclusion, we identified a novel variation of *GATA4* (c.958C>T, p.R320W) in familial ASD. The identified variant caused impaired biological function of the protein in vitro, suggesting that it is likely to play a role in the pathogenesis of ASD. Reasonable use of a variety of variant detection techniques will help us to identify pathogenic variants in CHD patients, especially in familial disease. Elucidation of the genetic basis of CHD has valuable clinical implications and will continue to expand our understanding of the pathogenesis of CHD.

## Acknowledgments

We sincerely thank all the patients and their family members for their enthusiasm and continued participation in this study. We also thank the clinicians and physicians for sending blood samples.

